# Conformational dynamics of electrostatically charged Intrinsically Disordered Proteins

**DOI:** 10.1101/2024.06.04.597329

**Authors:** Simon Schwarz, Subhartha Sarkar, Damir Sakhapov, Jörg Enderlein, Anja Sturm, Max Wardetzky

## Abstract

We experimentally and computationally study the dynamics of homogeneously electrically charged intrinsically disordered proteins (IDPs) in ionic solutions (aqueous solutions containing salt). Computationally, this is achieved by extending a recently introduced discrete elastic rod (DER) model to include electrostatic interactions. The respective model includes only three free parameters: bending stiffness, bond length, and hydrodynamic radius. These parameters were obtained by fitting the measured conformational dynamics of uncharged polypeptides as reported in a previous publication [8]. We compare our computational results with photo-electron transfer fluorescence correlation spectroscopy (PET-FCS) measurements of the conformational dynamics of the highly charged (-44 e^*−*^) intrinsically disordered protein Prothymosin α. We here report on an agreement between the loop-closing rates obtained from our computational model and the respective experimental measurements, with rate values significantly slower than those observed for uncharged IDPs.

## INTRODUCTION

Experimental evidence shows that more than 30% of all eukaryotic proteins are Intrinsically Disordered Proteins (IDPs), which do not form a folded three-dimensional structure [19]. These proteins play an important role in protein function and interaction [18].

From a physics point of view, IDPs can be considered as unstructured filaments driven by thermal Brownian motion, and their conformational dynamics has been modeled successfully with the discrete elastic rod (DER) model, see e.g. [9], as well as [1–3, 15, 16] for a mathematical derivation and treatment of the model. However, the simulations resulting from the DER model have thus far not taken into account any intramolecular interaction such as electrostatic forces caused by electric charges of an IDPs amino acids. This has not been an issue for glycine-serine repeat IDPs that were experimentally studied and modeled in [9] since in physiological buffer the constituent amino acids glycine and serine remain electrically neutral. However, if an IPD contains considerable amounts of the positively charged amino acids arginine and lysine or the negatively charged amino acids aspartate or glutamate, electrostatic interactions are no longer negligible.

In the present paper, we report on employing Photo-induced Electron-Transfer Fluorescence Correlation Spectroscopy (PET-FCS) to measure the loop-formation contact rates of the highly negatively charged IDP Prothymosin α (ProT-α) in aqueous buffer and in various salt solutions (Na^+^, K^+^, Ca^2+^, Mg^2+^) of varying concentration and with different quencher positions, see the section on experimental methods below for details.

Our measurements show that the charge distribution of the peptide has a decisive influence on the conformational dynamics. We correlate the measured dynamics with the conformational dynamics obtained from simulations based on our DER model including electrostatic interactions and show that our model is capable of approximately predicting the measured contact rates. Moreover, we find that even small changes of the charge distribution, e.g., resulting from inserting the quencher at a different position, lead to considerable variations in the contact rate.

Our simulations treat a physical time of more than 30 *µs*. On these time scales, we require efficient algorithms for computing Brownian motion on general Riemannian manifolds in order to simulate the thermal fluctations in the DER model. Here, we use a novel approach for efficiently sampling random walks based on *retractions*, i.e., approximations of the Riemannian exponential map, which has been analyzed in [17] and proven to converge to the correct dynamics.

## MATHEMATICAL MODEL

In order to model the dynamics of the Prot-α we follow the approach of [8, 9] and define a coarse grained model based on discrete elastic rods (DER), see [1–3], combined with additional Coloumb forces. The description of the model closely follows the presentation in [9].

### Discrete elastic rods

Originally designed as a structure-preserving version of smooth elastic rods including the delicate interplay between bending and twisting forces, the DER model was also found to be useful as a meso-scale model for proteins with bead positions **x**_1_, …, **x**_*n*+1_ connected by *inextensible* straight line segments **e**_1_, …, **e**_*n*_, see Figure 2.

**FIG. 1.**
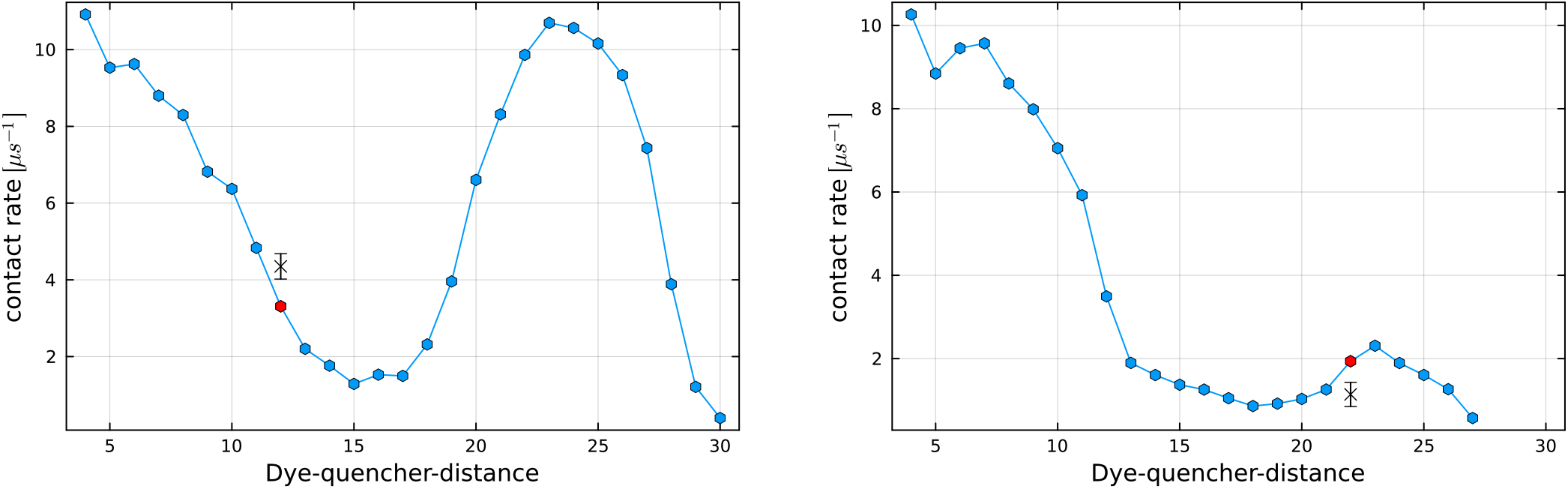
Simulated loop closing rate *k*+ for ProT-α modified by the insertion of a (neutrally charged) quencher at position 12 *(left)* and 22 *(right)*. The simulated closing rates of the respective quenchers are shown in red *(left:* 3.31 (*µs*)^*−*1^; *right:* 1.97 (*µs*)^*−*1^) while the experimentally measured rates are displayed in black *(left:* 4.35 ± 0.33 (*µs*)^*−*1^; *right:* 1.14 ± 0.29 (*µs*)^*−*1^). Notice that the slightly changed charge distribution resulting from replacing one positively charged amino acid at position 22 by the quencher has a tremendous impact on the loop closing rates.

**FIG. 2.**
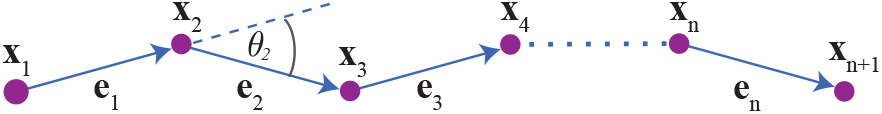
Polygonal line segments connecting (*n* + 1) point-like particles (*beads*).

The inextensibility constraint resembles strong intermolecular bonds – in the following, the bond length is fixed to *d*_0_ = 3.8 Å.

Thermal fluctuations are modeled as random motions of beads. Due to the inextensibility constraint this leads to Brownian motion (and more general diffusion processes) on the resulting configuration space – a sub-manifold of Euclidean space R^3(*n*+1)^. Here, we only consider a discretization of the bending energy within the DER model [2, 3], neglecting the (more elaborate) model for twist. This is justified by numerical experiments in [8], which show that the twist has only very little influence on contact rates.

As a discretization of the bending energy of the resulting polygonal space curve, one first considers the discrete dual lengths

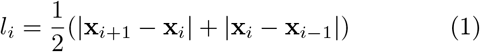

at vertex positions *i* and the discrete curvatures

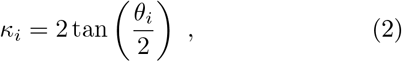

where *θ*_*i*_ = arccos (**e**_*i−*1_ **e**_*i*_) is the bending angle at vertex *i*. The discrete bending energy is then defined by

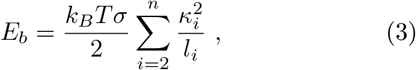

where the free scalar parameter *σ* is the *bending stiffness* of the elastic rod, *k*_*B*_ is the Boltzmann constant, and *T* denotes the temperature.

### Diffusion tensor

The dynamics of thermal fluctuations of proteins is strongly influenced by the diffusion tensor that accounts for hydrodynamic interactions. The diffusion tensor **D** is a symmetric, positive definite 3*N* × 3*N*-matrix, whose construction we here briefly sketch. For a more thorough derivation we refer to [8], which employs the Rotne-Prager approximation [12, 21]. We denote the hydrodynamic radius of the *i*-th bead by *a*_*i*_. In our case, all beads – except the fluorophore at position one – are equal. In particular, *a*_1_ = 6 Å and *a*_*i*_ = *a* for *i* = 2, …, *N*, where *a* is a free model parameter. The diagonal 3 × 3-blocks of **D** are then given by **D**_*ii*_ = *D*_0_1_3 × 3_, where *D*_0_ is a single bead’s scalar diffusion coefficient

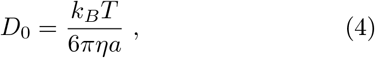

where *η* denotes the viscosity of the solution. Using the notation **x**_*ij*_ = **x**_*i*_ *−* **x**_*j*_, *x*_*ij*_ = |**x**_*ij*_|, and *a*_*ij*_ = *a*_*i*_ *− a*_*j*_, the other block matrices 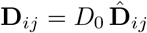 are defined as follows:

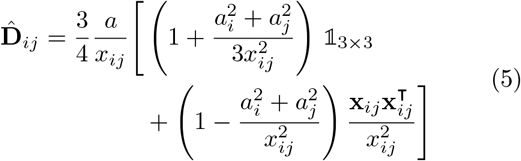

if *a*_*i*_ + *a*_*j*_ *< x*_*ij*_ (no hydrodynamical overlap), by

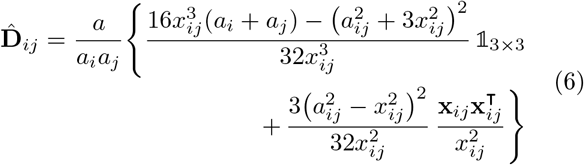

if |*a*_*ij*_|*< x*_*ij*_ *≤ a*_*i*_ + *a*_*j*_ (partial hydrodynamical overlap), and by

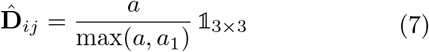

if *x*_*ij*_*≤* |*a*_*ij*_| (total hydrodynamical overlap).

*Equations of motion*. The motion of the beads **x**_*i*_, *i* = 1, …, *n*+1 is governed by a system of Langevin equations defined on the configuration space [5, 7, 9]. In particular,

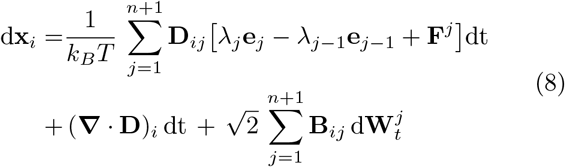

for all *i* = 1, …, *n* + 1, where (**W**_*t*_)_*t≥*0_ denotes the standard (*n* + 1)-dimensional Brownian motion, **D** = **BB**^⊺^ refers to the diffusion matrix from the previous paragraph (and its Cholesky decomposition), and the terms *λ*_*j*_**e**_*j*_ model tension forces with Lagrange multipliers *λ*_*j*_ determined by the geometric constraints

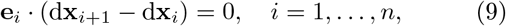

which correspond to linearizing the bond length constraints. Moreover, the force **F**^*j*^ is assembled by bending, Coloumb, and entropic force contributions:

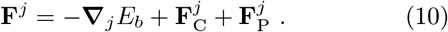

Here, *E*_*b*_ is the bending energy defined in Equation (3) and the Coulomb force is given by

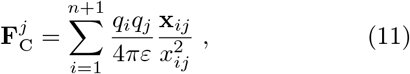

where *q*_*j*_ is the charge of bead **x**_*j*_ and *ε* the electric permittivity of the surrounding solution. For details on the conformation-dependent entropic force 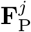 we refer to [8].

### Excluded volume

In order to prevent self-crossings of the chain, we use the <u>s</u>mallest possible excluded volume radius 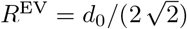, see [8], where *d*_0_ = 3.8 Å is the fixed bond length. If the distance between two beads gets smaller than this radius after integrating Equation (8) for some timestep, we reject the respective step and compute a new proposal. Notice that the excluded volume also prevents blow ups in the Coulomb force in Equation (11).

### SIMULATION AND DATA ANALYSIS

We conducted simulations of the conformational dynamics of ProT-α, for which we incorporated the electrostatic effects by including the Coulomb forces that result from the distribution of negative and positive charges along the polymer chain, see Figure 4. Notice that due to the insertion of a (neutrally charged) quencher at positions 12 and 22, the charge distribution of the polymer changes.

For the correct choice of the two free parameters of our simulation model, we rely on previous work in [9], in which they considered uncharged IDPs and derived the hydrodynamic radius *a* = 3.5 Å and bending stiffness *σ* = 1.83 as parameters. Simulations were conducted using projection-based (second order) retractions, see Section 2.2 of [17] for detail. The starting configuration of the polymer chain was generated from the appropriate Boltzmann distribution via an MCMC sampler, see [8]. Subsequently, the simulations were run for a physical time of 33 µ*s*, each integration step corresponding to about 220 fs. This time window was chosen in such a way that (at least for the measured dye-quencher distances) sufficiently many loop closing and opening events could be observed for a reliable estimate of the loop closing rate while at the same time keeping the needed computational effort feasible. We emphasize that this time scale is entirely out of reach for more detailed molecular dynamics (MD) simulations.

In Figure 1 we show the simulated loop closing rate of ProT-α in water (zero ionic concentration) as a function of the dye quencher distance. The charge distributions that we use in the simulations correspond to the charge distribution of ProT-α with a quencher inserted at position 12 *(left)* and 22 *(right)* respectively. Notice that the change of the polymer due to the insertion of the quencher has a significant impact on the loop closing rates from about bead 15 onwards. It is therefore crucial to adjust the polymer configuration to this change. The value obtained by our simulations are indeed a good approximation of the experimentally measured value, see again Figure 1 and Table I.

**TABLE 1.** The quencher was inserted at positions 12 and 22. In the table we compare the loop closing rates from the measurements and from the simulations using the DER model. In the displayed values of the experimental measurements we have already taken into account that only every third encounter leads to successful quenching, see [4, 20]. Thus, the measured rates have been multiplied by a factor of three.

| Quencher pos. | Measurement | Simulation |
| --- | --- | --- |
| 12 | $4.35 \pm 0.33 (\mu s)^{-1}$ | $3.31 (\mu s)^{-1}$ |
| 22 | $1.14 \pm 0.29 (\mu s)^{-1}$ | $1.97 (\mu s)^{-1}$ |

### EXPERIMENTAL METHODS AND RESULTS

For measuring the conformational dynamics of the IDP ProT-α, we have used Photo-induced Energy Transfer Fluorescence Correlation Spectroscopy (PET-FCS) [10, 11, 14]. In contrast to Förster Resonance Energy Transfer (FRET) spectroscopy, PET-FCS requires only one fluorescence label, the oxazine dye Atto655, which is efficiently quenched in its fluorescence when coming in direct contact with the amino acid tryptophan. This quenching is caused by an electron transfer from the excited state of the dye to the tryptophan, and is quasi hundred percent in the dye-tryptophan complex.

For measuring the conformational dynamics of a peptide, one covalently binds the dye to the C-terminus of the peptide and incorporates at one specific position one tryptophan into the peptide chain, which does otherwise not contain another tryptophan. By using fluorescence correlation spectroscopy (FCS) [6], one measures the quenching rate of the dye which shows up as a fast partial correlation decay in the time range of a few dozen nanoseconds to microseconds. Fortunately, the dye Atto655 shows hardly any significant triplet state kinetics [13] so that all nanosecond/microsecond correlation decay can be fully attributed to fluorescence quenching induced by direct contact formation between the dye and the tryptophan. By measurements on pure dye and tryptophan solutions [4, 20] it has been shown that, on average, every third dye-tryptophan contact leads to total fluorescence quenching. Thus, measuring the quenching rate between the dye and the tryptophan with FCS provides direct information about the loop closing rate of the peptide chain.

More precisely, FCS records the fluorescence intensity as a function of time (fluorescence time trace) and then calculates a correlation function *g*(*τ*) as

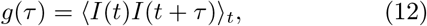

where *τ* is the correlation lag time, *I*(*t*) is the measured fluorescence intensity at time *t*, and the angular brackets denote averaging over time *t*. The loop formation kinetics (dye-tryptophan contact formation) is well described by a two-state system (open state = brightly fluorescent, and closed state = dark non-fluorescent) with exponentially distributed transition times between the two states (first order transition kinetics). In that case, the correlation function *g*(*τ*) takes the simple form:

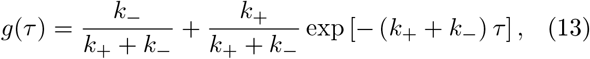

where *k*_+_ and *k*_*−*_ are the transition rates from the bright to the dark state and *vice versa*, i.e. the loop closing and loop opening rates, respectively. Thus, a PET-FCS measurement directly yields information about these rates *k*_+_ and *k*_*−*_. It should be noted that the off-rate *k*_*−*_ of peptide loop opening is strongly biased by the interaction energy between the dye Atto655 and the tryptophan, but the loop closing rate *k*_+_ is unbiased by this contact-pair energy and thus reliably reports about the peptide chain’s conformational dynamics. This is why the comparison of the experimental data with the simulation focuses on the loop closing rate *k*_+_, see Table I.

In the experiment we found that the high electrostatic charge of the protein chain ProT-α, see Figure 4, leads to strong electrostatic intra-chain repulsion and increases the apparent stiffness of the protein chain. This resulted overall in a smaller loop closing rate, in addition to the expected decrease of the loop closing rate as a function of dye-quencher distance as observed in [9]. As a result the loop closing rate at larger distances was too small to be observable in the experimental time frame.

We also measured the loop closing and loop opening rates for ProT-α in four different salt solutions (two monovalent salts NaCl and KCl, and two divalent salts CaCl_2_ and MgCl_2_) with fixed distance of 12 amino acids, see Figure 3. The idea behind this approach was that an increasing ionic strength would increasingly lead to electrostatic screening of the protein charges and should thus facilitate chain dynamics (modulating the overall chain bending stiffness). However, we found a linear slowing down of chain dynamics as a function of salt concentration solely due to the corresponding increase of viscosity that dominates any further screening effects as, for example, described by the Debye–Hückel theory.

**FIG. 3.**
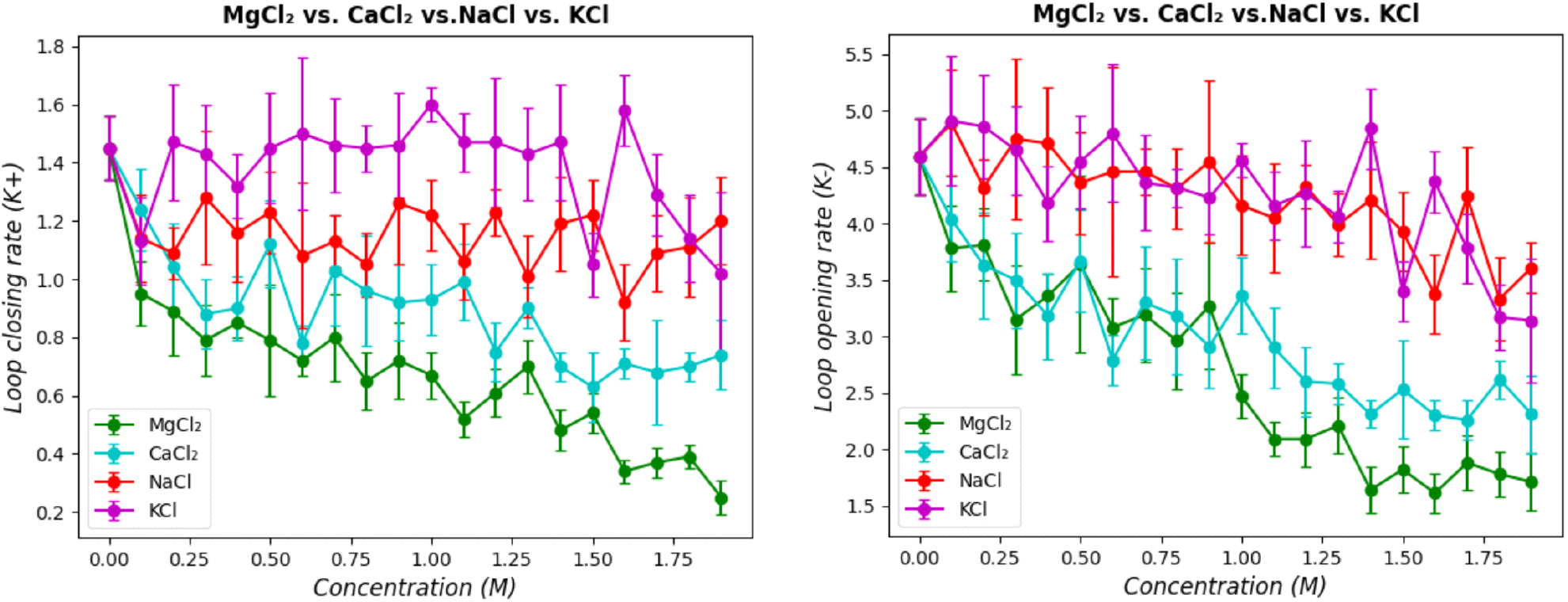
Results of Photo-electron-Transfer-Fluorescence-Correlation (PET-FCS) measurements of the opening and closing dynamics between two sites with 12 amino acid distance along a ProT-α protein as a function of ionic strength and salt composition. The left panels shows the values determined for the loop closing and the right panel for the loop opening rates in units of 1 µs. The figures summarize the results of 80 individual PET-FCS measurements.

**FIG. 4.**
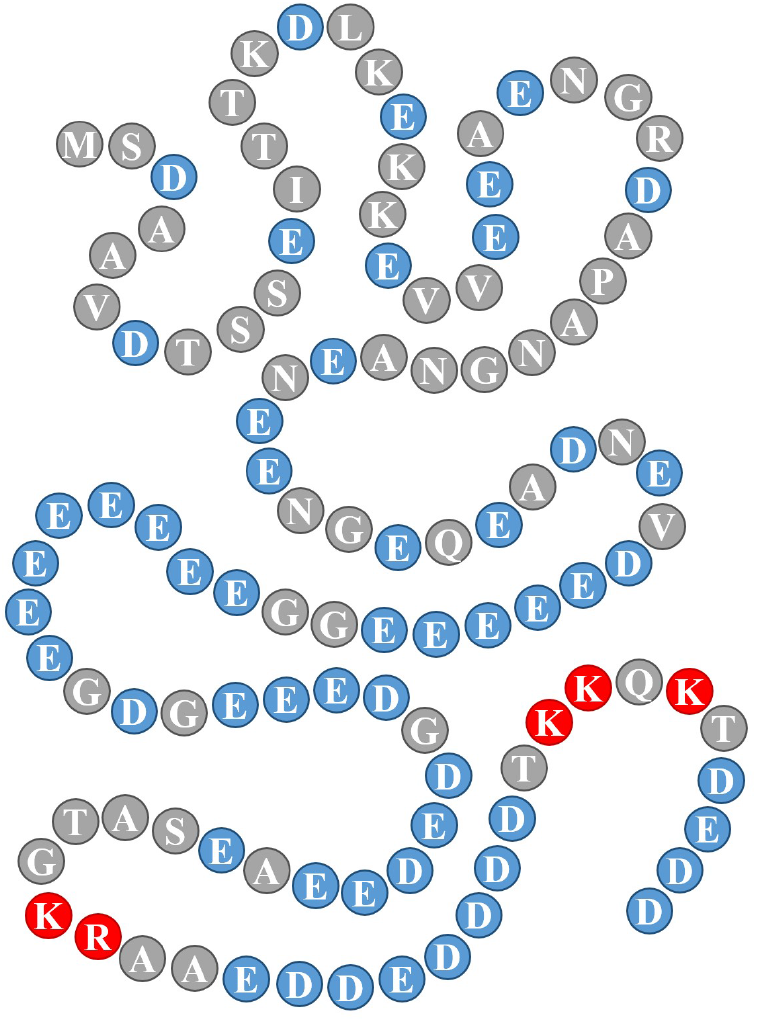
Amino acid sequence and distribution of charges along ProT-α. (Grey, blue and red circles are the neutral, negatively charged and positively charged amino acids, respectively.) The (neutrally charged) dye is attached to the end of the chain at the bottom right. Adding a quencher changes the structure of Prot-α depending on its position.

## CONCLUSION

We explored the conformational dynamics of the highly charged intrinsically disordered protein ProT-α. We combined experimental measurements using Photoinduced Electron-Transfer Fluorescence Correlation Spectroscopy (PET-FCS) with computational simulations based on an extended discrete elastic rod model.

Our experimental findings revealed that the electrostatic intra-chain repulsion due to high charge density in ProT-α notably increases the apparent stiffness of the protein chain, which in turn affects its conformational dynamics. Specifically, we observed a reduced loop closing rate that is sensitive to minor modifications of the charge distribution. We moreover show that salt solutions are not capable of reducing the electrostatic interactions, but rather slow down the chain dynamics even further due to a higher viscosity.

The computational simulations, incorporating Coulomb forces between charged amino acids, provided results that were in good agreement with the experimental data, validating the efficacy of the discrete elastic rod model in capturing the essential features of electrostatically driven conformational changes.

## Acknowledgement

Financial support by the Deutsche Forschungsgemeinschaft is gratefully acknowledged (SFB 1456, project A02).

